# A non-canonical cGAS-STING pathway drives cellular and organismal aging

**DOI:** 10.1101/2025.04.03.645994

**Authors:** Rafael Cancado de Faria, Lilian Silva, Barbara Teodoro-Castro, Kyle S. McCommis, Elena V. Shashkova, Susana Gonzalo

## Abstract

Accumulation of cytosolic DNA has emerged as a hallmark of aging, inducing sterile inflammation. STING (Stimulator of Interferon Genes) protein translates the sensing of cytosolic DNA by cGAS (cyclic-GMP-AMP synthase) into an inflammatory response. However, the molecular mechanisms whereby cytosolic DNA-induced cGAS-STING pathway leads to aging remain poorly understood. We show that STING does not follow the canonical pathway of activation in human fibroblasts passaged (aging) in culture, senescent fibroblasts, or progeria fibroblasts (from Hutchinson Gilford Progeria Syndrome patients). Despite cytosolic DNA buildup, features of the canonical cGAS-STING pathway like increased cGAMP production, STING phosphorylation, and STING trafficking to perinuclear compartment are not observed in progeria/senescent/aging fibroblasts. Instead, STING localizes at endoplasmic reticulum, nuclear envelope, and chromatin. Despite the non-conventional STING behavior, aging/senescent/progeria cells activate inflammatory programs such as the senescence-associated secretory phenotype (SASP) and the interferon (IFN) response, in a cGAS and STING-dependent manner, revealing a non-canonical pathway in aging. Importantly, progeria/aging/senescent cells are hindered in their ability to activate the canonical cGAS-STING pathway with synthetic DNA, compared to young cells. This deficiency is rescued by activating vitamin D receptor signaling, unveiling new mechanisms regulating the cGAS-STING pathway in aging. Significantly, in HGPS, inhibition of the non-canonical cGAS-STING pathway ameliorates cellular hallmarks of aging, reduces tissue degeneration, and extends the lifespan of progeria mice. Our study reveals that a new feature of aging is the progressively reduced ability to activate the canonical cGAS-STING pathway in response to cytosolic DNA, triggering instead a non-canonical pathway that drives senescence/aging phenotypes.

**Significance Statement:** Our study provides novel insights into the mechanisms driving sterile inflammation in aging and progeria. We reveal a previously unrecognized characteristic of aging cells: the progressive loss of ability to activate the canonical response to foreign or self-DNA at the cytoplasm. Instead, aging, senescent, and progeria cells activate inflammatory programs via a non-conventional pathway driven by cGAS and the adaptor protein STING. Importantly, pharmacological inhibition of the non-canonical cGAS-STING pathway ameliorates cellular, tissue and organismal decline in a devastating accelerated aging disease (Hutchinson Gilford Progeria Syndrome), highlighting it as a promising therapeutic target for age-related pathologies.

## Introduction

Presence of self-DNA in the cytosol is a hallmark of aging. Cytosolic DNA accumulates upon nuclear or mitochondrial damage, replication stress, reduced cytoplasmic nucleases, derepression of retrotransposons or transposable elements, and compromised autophagy (1-7). Cytosolic DNA is detected by many cellular sensors (8) and the best-studied is cGAS (9-13). Once activated by dsDNA or DNA:RNA hybrids, it synthesizes cGAMP and triggers the canonical STING pathway (14-16). Other sensors such as IFI16 cooperate with cGAS to activate STING (17) but can also activate STING independent of cGAS (18), yielding an entirely different transcriptional profile. These observations suggest that STING differs in its response depending on the activation mode.

STING (TMEM173), is localized at the endoplasmic reticulum (ER) (19) and nuclear envelope (NE) in the resting state (20). Upon cGAMP binding, STING undergoes conformational changes that result in its oligomerization (21, 22), its trafficking to a perinuclear compartment (PNC) (23) formed primarily by Golgi and vesicles/endosomes, while undergoing multiple post-translational modifications (PTMs), including phosphorylation, palmitoylation, and ubiquitylation (24, 25). At the PNC, STING forms a complex with TANK-binding kinase 1 (TBK1) and IFN-regulatory factor 3 (IRF3), which results in IRF3 phosphorylation and nuclear translocation (26, 27). STING also activates NFκB, inducing its nuclear translocation, and both IRF3 and NFκB upregulate expression of IFNs and inflammatory cytokines (9). IFNα/β, via IFNAR receptors, activate the JAK-STAT pathway that induces expression of IFN-stimulated genes (ISGs). This sterile inflammation and IFN response are linked to activation of the SASP (5, 28, 29), and often seen in aging-related conditions (30-32), including cardiovascular disease, cancer, and premature aging diseases (33, 34). Besides the role of STING in inflammation, it also regulates autophagy and cell death (35-37). Thus, STING is in the spotlight as a promising target in aging-related diseases (38-40).

HGPS is a premature aging disease caused by *LMNA* gene mutation and expression of a truncated lamin A protein named progerin, which triggers most hallmarks of aging: DNA damage, telomere dysfunction, loss of heterochromatin, mitochondrial dysfunction, loss of proteostasis, and early senescence. We reported that progerin causes replication stress (RS), cytosolic DNA buildup, and a sterile inflammation cascade that includes cytokines characteristic of SASP, and ISGs (34, 41). However, the molecular mechanisms whereby cytosolic DNA drives SASP and IFN response in progeria and in normal aging/senescence remain poorly understood.

Here we show that STING drives this sterile inflammation/IFN response in HGPS patients’ fibroblasts and aging fibroblasts (passaged until entering replicative senescence) via a non-canonical pathway. Despite cytosolic DNA accumulation, progeria, aged (late passage), and senescent fibroblasts (induced by ionizing radiation) do not activate the canonical cGAS-STING pathway, characterized by increased cGAMP production, phosphorylation of STING, and trafficking of STING to PNC. Instead, STING localizes at the ER and the nucleus, primarily at the NE and bound to chromatin. Nevertheless, these cells activate inflammatory programs, such as SASP and IFN response, in a cGAS- and STING-dependent manner, suggesting the existence of a non-canonical cGAS-STING pathway in aging. Moreover, forced activation of the canonical pathway with synthetic dsDNA in progeria, aged, and senescent cells leads to lower cGAMP production and reduced STING phosphorylation and trafficking to PNC compared to young cells (early passage). Interestingly, rejuvenating progeria cells with calcitriol (activates vitamin D receptor signaling) rescues the canonical pathway in response to synthetic DNA, unveiling a novel mechanism regulating the canonical cGAS-STING pathway in aging. Inhibition/depletion of STING reduces SASP and the IFN response and improves cellular fitness in progeria cells. Targeting STING increases the lifespan of progeria mice (*Lmna*^*G609G/G609G*^), with remarkable improvements in white adipose tissue (WAT) and vascular smooth muscle cells (VSMCs) in the aorta; two tissues with high degeneration in HGPS. These studies unveil a different behavior of STING in response to cytosolic DNA in old vs young cells, and a key role of the non-canonical cGAS-STING pathway driving aging phenotypes. We also provide strong evidence for targeting STING as a therapeutic strategy for diverse pathologies associated with physiological aging and progerias.

## Results

### Cytosolic DNA buildup in progeria cells does not activate the canonical cGAS-STING pathway

We previously found that progerin-expressing fibroblasts exhibit genomic instability, nuclear fragility, and accumulation of cytosolic DNA (41). We also found increased levels of cGAS and STING proteins, and upregulation of a robust innate immune/IFN-like response (42). To test whether progerin causes activation of the canonical cGAS-STING pathway, we used a doxycycline-inducible GFP-progerin expression system, generated by lentiviral transduction of human derived fibroblasts (HDF-*tet*^*on*^-GFP-progerin) (43). After 4-8 days, GFP-progerin expression triggers aging phenotypes characteristic of HGPS fibroblasts (41, 42). We find that despite the accumulation of DNA damage and cytosolic DNA (41) (**Supplemental Figure 1A-C**), expression of progerin (doxy for 2-16 days) does not lead to a detectable increase in the production of cGAMP (monitored by ELISA) (**Figure 1A**, 8-days doxy). In contrast, transfection of poly(dA:dT), synthetic DNA that activates the canonical cGAS-STING pathway, leads to a robust increase in cGAMP levels in HDF control. Importantly, cGAMP synthesis in response to poly(dA:dT) is significantly reduced in progerin-expressing cells, suggesting an inhibitory effect on cGAS enzymatic activity.

**Figure 1.**
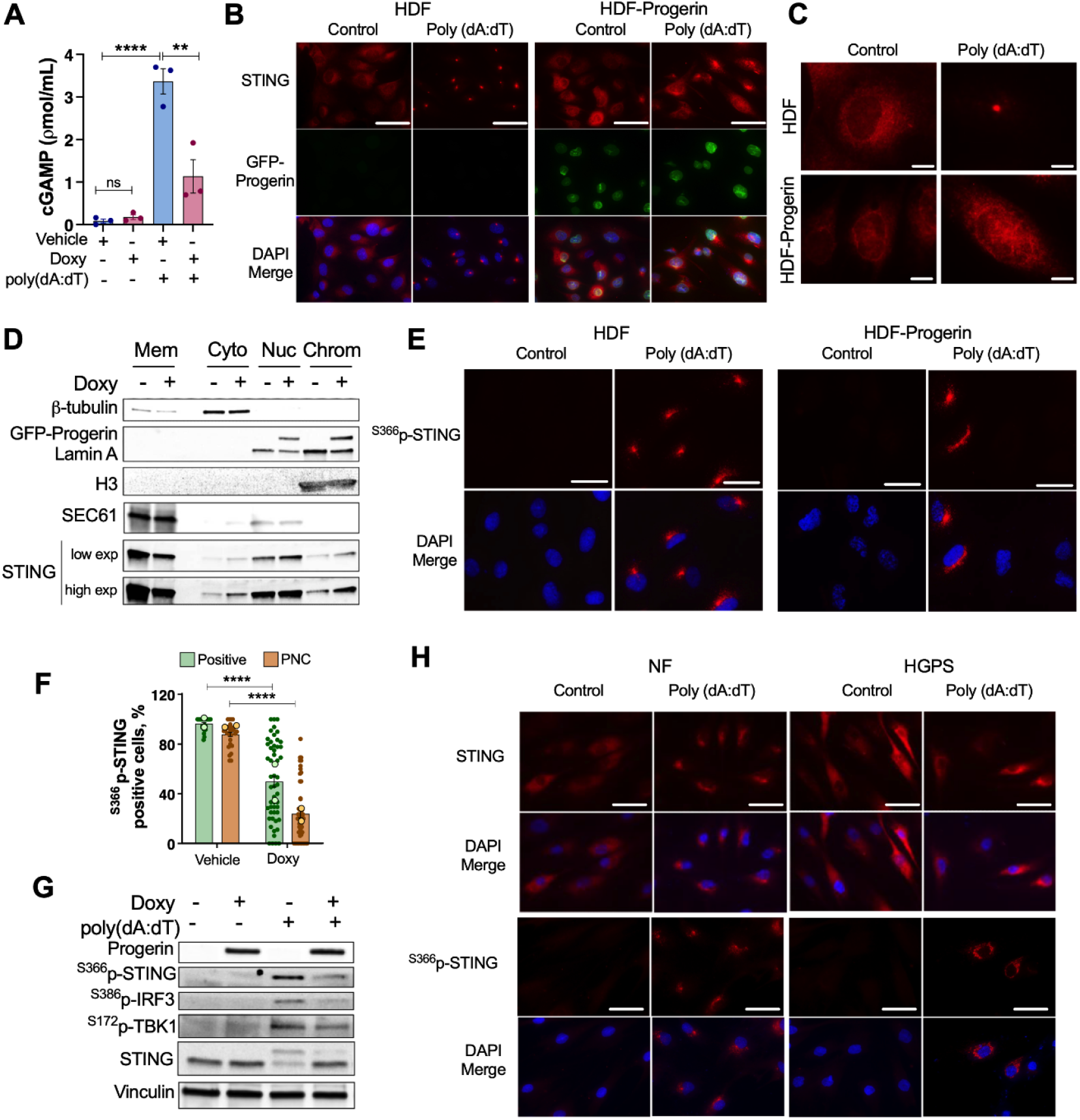
Progeria cells are impaired in the ability to activate the canonical cGAS-STING pathway. (**A**) Human derived fibroblasts (HDF) induced to express GFP-progerin (HDF-tet^on^-GFP-progerin) with doxycycline (0.5μg/mL) for 8 days (DMSO as vehicle). cGAMP synthesis measured by ELISA. cGAMP concentration was also assessed 4 hrs after transfection of 1μg/mL poly(dA:dT). Data represent average±SEM of 3 biological repeats. (**B**) Immunofluorescence (IF) with STING antibody (red) in the same cells as in (A). GFP-progerin is shown in green and DNA staining (DAPI) in blue. Scale bars: 50μM. (**C**) Higher magnification images showing STING localization. Scale bars: 10μm. (**D**) Subcellular fractionation to monitor STING presence at membranes, cytoplasm, nucleus, and chromatin fractions in HDF-tet^on^-GFP-progerin cells. β-tubulin used as cytoplasm fraction control, SEC61 as ER marker, Lamin A as nuclear fraction control, and H3 as chromatin marker. Mem: membrane fraction; Cyto: cytoplasmic soluble fraction; Nuc: nuclear fraction including nuclear envelope; Chrom: chromatin fraction. (**E**) IF with ^S366^p-STING antibody (red) and DAPI (blue) in HDF-tet^on^-GFP-progerin. Scale bars: 50μm. (**F**) Percentage of ^S366^p-STING positive cells or cells with ^S366^p-STING at the PNC. Mean of each of two independent experiments is shown (light circles), as well as mean±SEM of all the image fields of both experiments (∼25 image fields/experiment and >200 cells/experiment). Statistical significance via two-way ANOVA. (**G**) Immunoblot of HDF-teton-GFP-progerin transfected with poly(dA:dT) shows phosphorylation of STING, TBK1, and IRF3 in control HDF that is reduced upon progerin expression. Vinculin used as loading control. (**H**) IF showing STING and ^S366^p-STING localization ± poly(dA:dT) stimulation in NF or HGPS fibroblasts. Scale bars: 50μm. DAPI (blue) used for nuclear staining.

Other hallmarks of the canonical cGAS-STING pathway are STING trafficking out of the ER, concentrating at cytoplasmic puncta known as PNC, and STING phosphorylation on Ser366. We monitored these features by immunofluorescence with STING and ^S366^p-STING antibodies in HDF-*tet*^*on*^-GFP-progerin model. STING localizes at the ER and NE in resting control HDF, and activation of the canonical cGAS-STING pathway with poly(dA:dT) leads to STING translocation to PNC (**Figure 1B**). Expression of progerin does not lead to STING trafficking to PNC. Rather, STING presence at the nucleus (NE and nuclear interior) is increased. Even after forced activation of the canonical pathway with poly(dA:dT), progerin-expressing cells show deficiencies in STING trafficking to PNC, with many cells accumulating STING in the nucleus instead. Differences in STING localization are clearly observed in **Figure 1C** (single cells at higher magnification). Consistently, subcellular fractionation reveals that progerin expression increases STING levels in the nucleus and bound to chromatin (**Figure 1D**, densitometry in **Supplemental Figure 1D**). We also found that poly(dA:dT) triggers STING phosphorylation (^S366^p-STING) in control HDF, which localizes to PNC (**Fig 1E**). Progerin-expressing HDFs show a diminished response to poly(dA:dT), with decreased percentage of cells featuring ^S366^p-STING or localization of ^S366^p-STING at PNC, compared to control HDF (**Figure 1F**). We did not find however any differences in STING dimerization or high-order oligomerization upon activation by poly(dA:dT) in control vs progeria cells (**Supplemental Figure 1E**), suggesting that alterations of STING behavior occur independent of oligomerization. Thus, progerin expression hinders the ability of HDF to activate the canonical cGAS-STING pathway by synthetic DNA. These findings were confirmed by immunoblotting phosphorylated forms of STING, IRF3, and TBK1 (**Figure 1G**), which form a ternary complex at PNC that activates inflammatory genes as part of the canonical cGAS-STING pathway (8, 13, 30). We find phosphorylation of the complex in poly(dA:dT)-treated HDF but minimal upon progerin expression (up to 8 days). Activation of the canonical pathway (^S366^p-STING, ^S386^p-IRF3, and ^S172^p-TBK1) by poly(dA:dT) is reduced in progerin-expressing cells (**Figure 1G**, densitometry in **Supplemental Figure 1F**), consistent with the low cGAMP levels and reduced STING translocation to PNC. Also, unphosphorylated STING (lower band) is significantly reduced in poly(dA:dT)-treated HDF, consistent with its canonical activation (phosphorylation) and consequent degradation. In contrast, unphosphorylated STING remains high in progerin-expressing cells, consistent with the altered STING behavior in these cells.

Next, we monitored activation of the canonical cGAS-STING pathway in HGPS patients-derived fibroblasts, compared to normal fibroblasts (NF) from patients’ parents, by immunofluorescence. We did not find STING phosphorylation or localization at PNC in HGPS cells (**Figure 1H**). Transfection with poly(dA:dT) leads to STING phosphorylation and trafficking to PNC in nearly all NF, but a much-reduced number of HGPS cells show these features. Consistently, we did not detect increased production of cGAMP in HGPS cells by ELISA (see Figure 2 below). These data indicate that progerin does not activate the canonical cGAS-STING pathway characterized by elevated cGAMP synthesis and STING phosphorylation and trafficking to PNC, despite the accumulation of cytosolic DNA and the activation of SASP and the IFN response (42). Importantly, expression of progerin (HGPS and HDF-GFP-progerin) hinders the ability of cells to activate the canonical cGAS-STING pathway.

**Figure 2.**
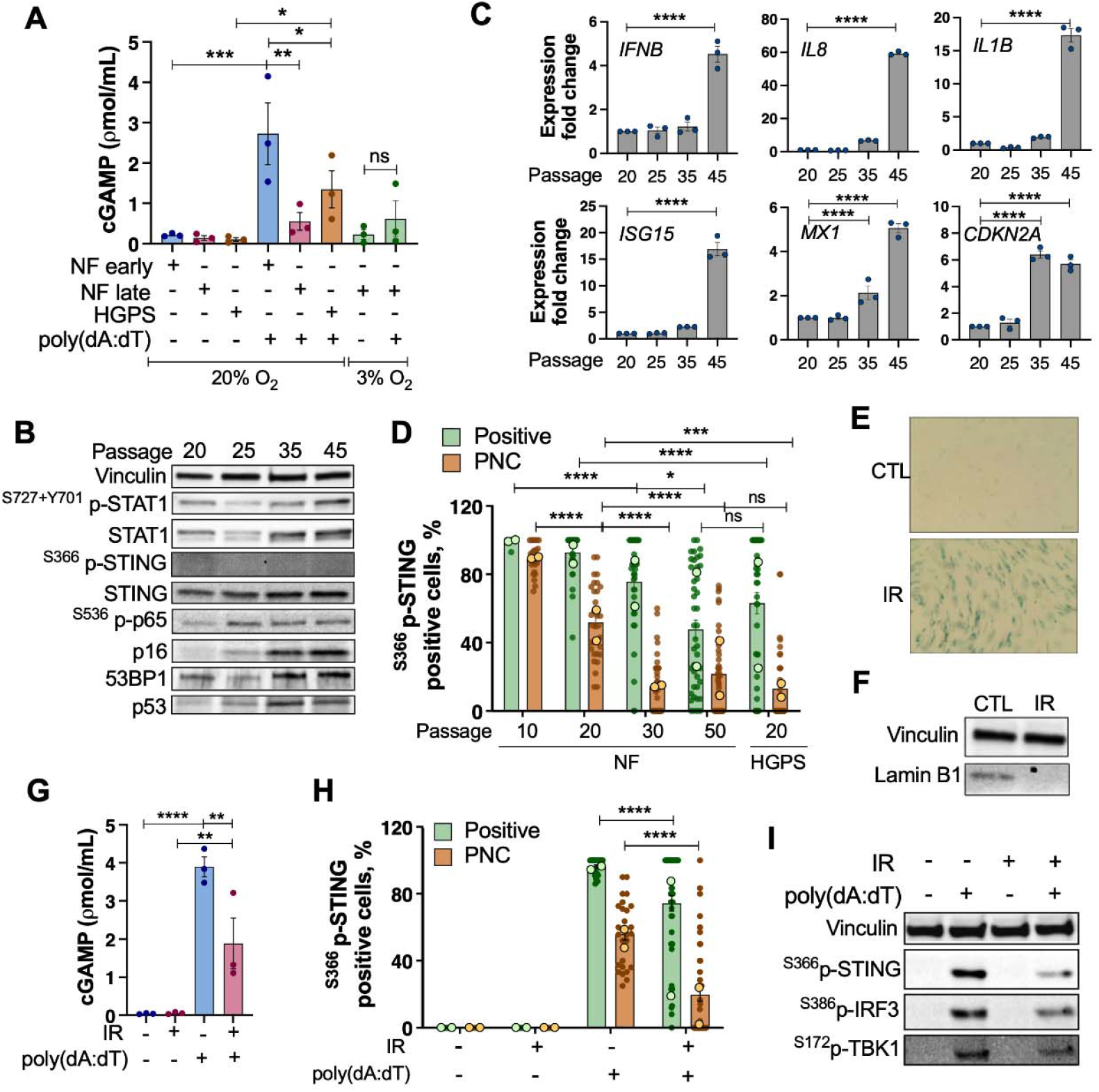
Canonical cGAS-STING pathway is hindered during passage of cells in culture and in DNA damage-induced senescent cells. (**A**) cGAMP production by ELISA in NF of early passage<20 and late passage>32, and HGPS patients (passage<16). cGAMP also measured after transfection with 1μg/ml poly(dA:dT) for 4h, and in cells cultured under 20% or 3% oxygen. Data represent average±SEM of 3 biological repeats. (**B**) Immunoblots of inflammation markers upregulated in NF during passage (STAT1, ^S727+Y701^p-STAT1, STING, and ^S536^p-p65), and other markers of senescence (p16, p53 and 53BP1). Not detected ^S366^p-STING at any passage. Vinculin was loading control. (**C**) qRT-PCR of SASP transcripts IFNB, IL8 and IL1B; senescence marker CDKN2A; and ISGs (ISG15 and MX1). Graphs show expression fold change of triplicates from a representative experiment from at least two biological repeats. (**D**) Percentage of ^S366^p-STING positive cells or cells with ^S366^p-STING at PNC, calculated as in Figure 1F. Mean of each of experiment (light circles) and average±SEM of all the image fields of both experiments (>200 cells/experiment). Statistical significance by two-way ANOVA. (**E**) NF early passage (P20) irradiated (20 Gy) and cultured for 3 weeks. β-galactosidase activity shows positivity in growth-arrested irradiated cells. (**F**) Lamin B1 immunoblot from the same cells as in (E). Vinculin was loading control. (**G**) cGAMP levels by ELISA. NF (P20) and senescent NF (P20+IR+3 weeks) were transfected with poly(dA:dT) to activate the canonical cGAS-STING pathway. Graph shows mean±SEM of 3 biological repeats. (**H**) Percentage of S366p-STING positive cells or cells with ^S366^p-STING at the PNC, in same cells as in (E). Average of two biological repeats (n>150 cells/experiment). Statistical analysis via two-way ANOVA. (**I**) Immunoblot of NF early passage control and senescent (3 weeks after IR) that were transfected with poly(dA:dT); phosphorylation of STING, TBK1, and IRF3 was monitored. Vinculin was loading control.

### Aging and senescence hinder the canonical cGAS-STING pathway

Given that cGAS and STING have been shown to be indispensable for senescence (44, 45) and crucial for SASP activation by NFκB, we determined if primary human fibroblasts activate the canonical cGAS-STING pathway during replicative senescence, or ionizing radiation (IR)-induced senescence. We found no differences in the levels of cGAMP between human primary fibroblasts of early and late passage (**Figure 2A**). At all passages, poly(dA:dT) induced cGAMP synthesis, but this was significantly lower in late passage and HGPS fibroblasts, indicating that cGAS activity in response to synthetic DNA decreases in contexts of aging. This deficiency is not due to oxidative stress accumulating in culture, because we obtained the same results in cells cultured in low O_2_ (3%) conditions.

We confirmed that cells acquire senescence markers during passage in culture, such as high p16 expression, increased markers of DNA damage p53 and 53BP1, NFκB activation by increased p65 phosphorylation at Ser536 (^S536^p-p65), and increased levels of STAT1 and phosphorylated active STAT1 (^S727+Y701^p-STAT1) (**Figure 2B**). In contrast, we did not detect STING phosphorylation at any point during the passage of fibroblasts in culture. Moreover, we found increased transcripts of SASP genes (*IFNB, IL8, IL1B*), ISGs (*ISG15 and MX1*) and p16 (*CDKN2A*) (**Figure 2C**), consistent with activation of SASP and ISGs as cells approach replicative senescence. Importantly, NF of early passage (P10-20) can phosphorylate and transport STING to the PNC upon activation of the canonical pathway with poly(dA:dT) (**Supplemental Figure 2A**). However, as cells age in culture (NF late passage P30-50), they exhibit a reduced ability to phosphorylate STING (^S366^p-STING) and transport STING to the PNC (**Figure 2D**). These data demonstrate that STING behavior is altered during aging, and especially as cells enter replicative senescence, mirroring progeria cells.

Next, we determined whether early passage fibroblasts induced to undergo senescence exhibit the same altered behavior of STING as aged and progeria cells. NF at passage 20 were treated with ionizing radiation (20 Gy) and grown in culture changing the media regularly for three weeks; a point at which they entered senescence, as shown by increased β-gal activity (**Figure 2E**), loss of lamin B1 (**Figure 2F**), and upregulation of cytokines typical of SASP (*IL8, IL1B, IFNB*), and ISGs (*ISG15, MX1, STAT1, IFIT3*) (**Supplemental Figure 2B**). Note that the upregulation of cytokines and ISGs in early passage fibroblasts induced to undergo senescence mirrors that of late passage fibroblasts. Late passage (>50) and senescent fibroblasts (IR-induced) do show phosphorylation and nuclear translocation of p65 subunit of NFκB (**Supplemental Figure 2C**), another feature of senescent cells. However, these cells do not show elevated cGAMP levels (**Figure 3G**), STING phosphorylation or trafficking to PNC (**Figure 3H**). When transfected with poly(dA:dT), they also show deficiencies in the activation of the canonical pathway, with reduced cGAMP levels (**Figure 3G**), and decreased STING phosphorylation and PNC trafficking (**Figure 3H**). Moreover, IR-induced senescent cells show deficiencies in the phosphorylation of the STING/IRF3/TBK1 ternary complex in response to synthetic DNA, compared to non-senescent cells (**Figure 3I** and **Supplemental Figure 2D**).

**Figure 3.**
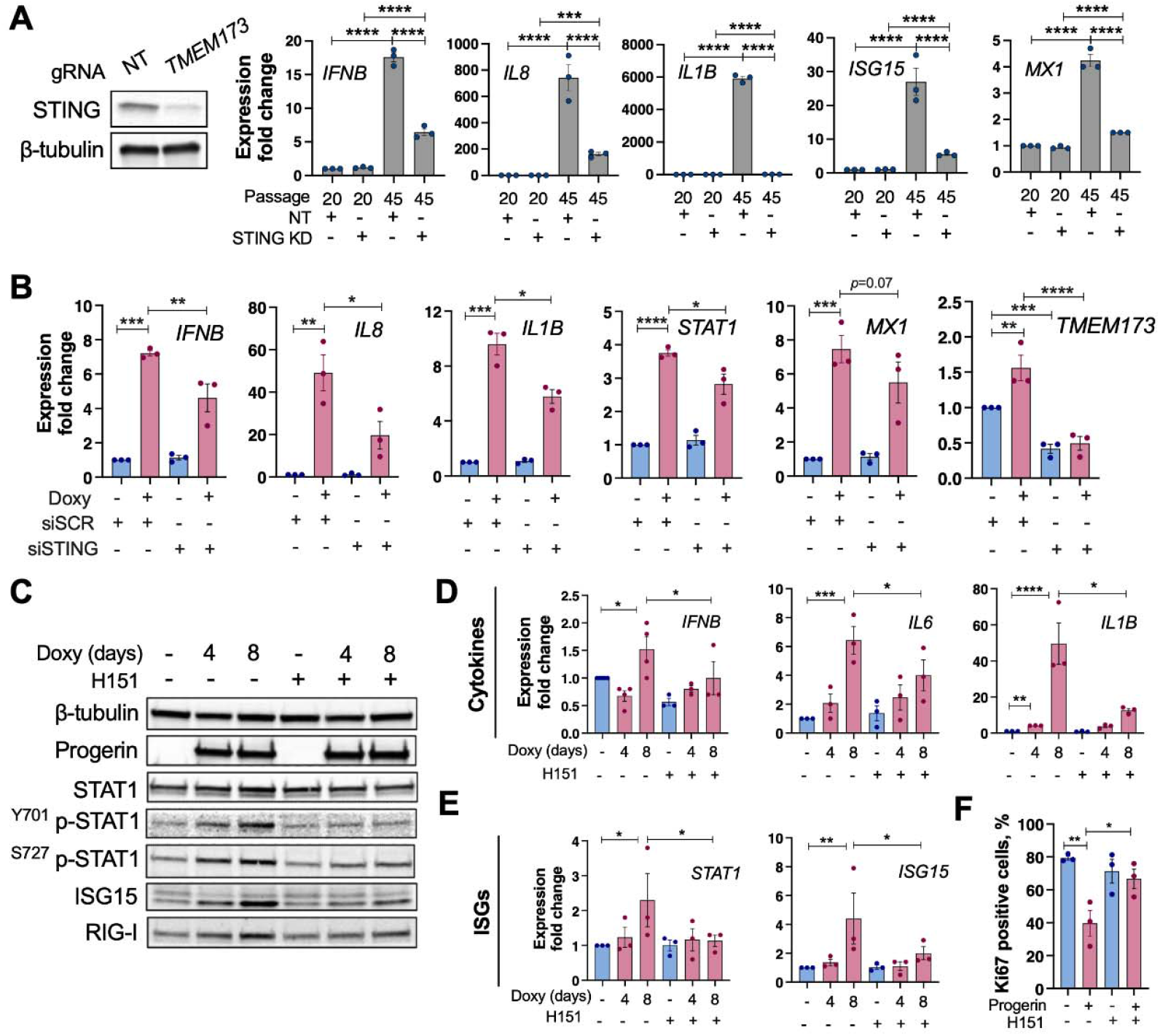
STING is required for SASP and IFN response in aging/progeria cells. (**A**) Immunoblots show depletion of STING in NF using gRNA/CRISPR/Cas9. Graphs show transcript levels of inflammatory cytokines (IFNB, IL8, IL1B) and ISGs (ISG15, MX1) upon STING depletion in NF of early (P20) and late passage (P45). One representative experiment out of two independent biological repeats shown. Note how the increase in all inflammatory markers in late passage NF is reduced by STING depletion. (**B**) HDF-tet^on^-GFP-progerin were transfected with siRNA targeting STING, and inflammatory cytokines (IFNB, IL8, IL1B) and ISGs (STAT1, MX1) monitored by qRT-PCR. Note how the increased expression of inflammatory genes is ameliorated by STING depletion. (**C**) HDF-tet^on^-GFP-progerin induced to express progerin for 4-8 days were treated with vehicle (DMSO) or H151 (STING inhibitor) during the 4-8 days (0.5μM H151). Representative blots from three independent experiments showing markers of sterile inflammation/IFN response: STAT1, p-STAT1, RIG-I and ISG15. β-tubulin used as loading control. (**D-E**) qRT-PCR of inflammatory cytokines (IFNB, IL6 and IL1B) and ISGs (STAT1 and MX1) in HDF-tet^on^-GFP-progerin expressing progerin for 6 days and treated with H151 or vehicle. Graph shows average±SEM of three independent experiments. (**F**) Percentage of Ki67 positive cells (by IF) for proliferative capacity assessment in HDF-tet^on^-GFP-progerin cells. Graph shows average±SEM of three independent experiments.

These data demonstrate that normal fibroblasts undergoing replicative senescence or DNA damage-induced senescence do not activate the canonical cGAS-STING pathway. In addition, they are hindered in their ability to activate the pathway upon transfection of synthetic DNA, which in early passage (young) cells activates the entire population of cells.

### STING is required for sterile inflammation in senescent and progeria cells

To determine if despite its altered behavior, STING is responsible for mediating SASP and IFN response in aging cells, we generated STING-deficient early and late passage human fibroblasts via gRNA/CRISPR/Cas9 system (NT=non-targeted gRNA; TMEM173=STING targeted gRNA). (**Figure 3A**). We find that the increase in SASP genes (*IFNB, IL8, IL1B*) and ISGs (*ISG15, MX1*) with passage of fibroblasts in culture is reduced by STING depletion (**Figure 3A**). To determine whether SASP and IFN pathways are regulated by STING in progerin-expressing cells (HDF-*tet*^*on*^-GFP progerin), we performed transfection with siRNAs targeting STING. As shown in **Figure 3B**, expression of progerin for six days upregulates inflammatory cytokines (*IL8, IL1B*, and *IFNB*) and ISGs (*STAT1* and *MX1*), which are reduced by STING (*TMEM173)* depletion. Thus, despite its altered behavior, STING plays a role in sterile inflammation in aging cells. Our data indicate that progeria/aged/senescent cells are hindered in their ability to activate the canonical cGAS-STING pathway, triggering SASP and IFN response nevertheless, via a non-canonical pathway.

To determine how the non-canonical pathway contributes to cellular aging phenotypes, we used a pharmacological approach with progeria as a model. First, we tested the impact of inhibiting STING with the compound H151 on aging phenotypes such as sterile inflammation and proliferation defects in HDF-*tet*^*on*^-GFP-progerin. H151 is a potent, irreversible and selective small molecule inhibitor of STING (46). We treated HDF with H151 two days prior and during the induction of progerin expression. Cells collected after 4-8 days show activation of STAT1-mediated IFN-response, with phosphorylation of STAT1 at Tyr701 and Ser727, and upregulation of ISGs such as ISG15, RIG-I and STAT1, by immunoblotting (**Figure 3C**, densitometry in **Supplemental Figure 3A**). Interestingly, H151 suppresses this response. Similarly, transcript levels of inflammatory cytokines *IFNB, IL6* and *IL1B* (**Figure 3D**) and ISGs *STAT1* and *ISG15* (**Figure 3E**) are reduced by H151 (especially at day 8). In addition, the decrease in proliferative capacity (Ki67 positivity) in progerin-expressing cells is rescued by H151 treatment (**Figure 3F** and **Supplemental Figure 3B**). Thus, inhibition of the non-canonical STING pathway in a progeria cellular model reduces inflammatory pathways and improves proliferation, supporting the role of STING in cellular aging hallmarks.

### cGAS is required for sterile inflammation in progeria

Given the important role of STING in sterile inflammation, we next determined whether cGAS, and its enzymatic activity, participates in the inflammatory response in progeria cells. Depletion of cGAS ameliorates the STAT1-mediated IFN-response, with reduced STAT1 phosphorylation, and downregulation of ISGs such as ISG15 and STAT1 (**Figure 4A** and **Supplemental Figure 4A**). Similarly, transcript levels of *IFNB, STAT1, ISG15*, and *MX1* (**Figure 4B**) are reduced in cGAS-depleted progeria cells. In addition, treatment with an inhibitor of cGAS (G140) suppresses the STAT1-mediated IFN response (**Supplemental Figure 4B**) in progeria cells. Note how two concentrations of G140 reduce the levels of STAT1, ISG15, and phosphorylated STAT1. We also find reduced transcripts levels for *IFNB, STAT1, ISG15*, and *MX1* upon treatment with G140 (**Supplemental Figure 4C**). These data indicate that despite the low levels of cGAMP observed by ELISA, cGAS is necessary to ensure the IFN response in progeria cells. Our results are in concordance with other studies demonstrating the critical role of cGAS in senescence (2, 6, 7, 28, 29, 44). It is possible that progerin causes a modest increase in cGAMP, which does not reach the threshold of detection by ELISA, but it is sufficient to activate the non-canonical STING pathway. We can conclude that cGAS plays a key role in the activation of sterile inflammation in progerin-expressing cells, via a mechanism that differs from the canonical pathway.

**Figure 4:**
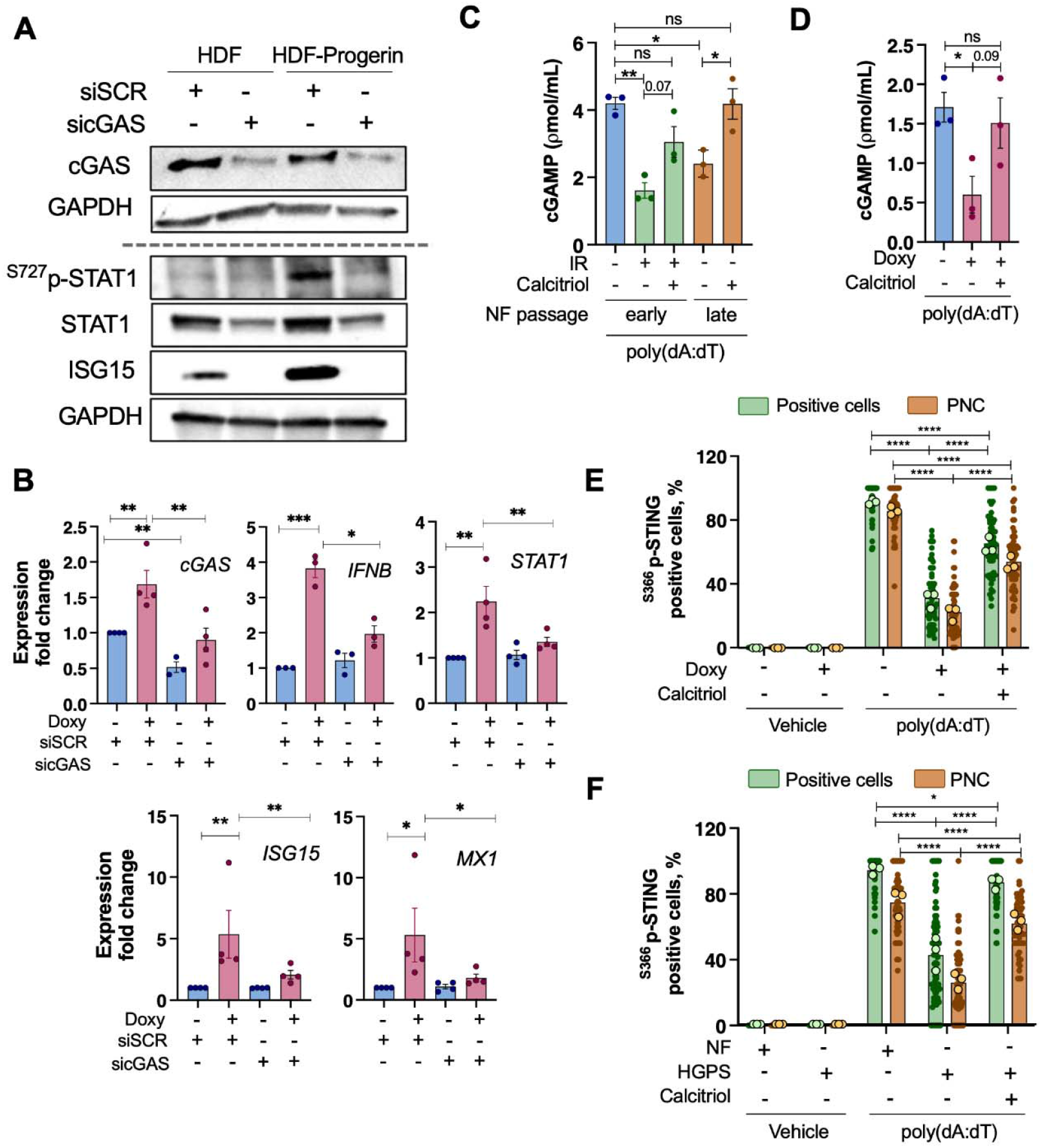
cGAS and calcitriol regulate the canonical and non-canonical cGAS-STING pathways. **(A)** HDF-tet^on^-GFP-progerin expressing progerin for 8 days (doxy) were transfected with siRNA targeting cGAS or siRNA control. Immunoblots show decrease in cGAS levels. Cell lysis protocol was modified to detect cGAS (see Methods). Immunoblots also show decreased STAT1, ^S727^p-STAT1, and ISG15. **(B)** qRT-PCR in the same cells as in (A) to monitor cGAS depletion and expression of inflammatory markers (IFNB, STAT1, ISG15, and MX1). **(C)** cGAMP levels by ELISA in IR-induced senescent cells, and in early and late passage NF transfected with 1μg/mL poly(dA:dT). Cells were incubated with calcitriol (100nM) where indicated. Data represent average±SEM of 3 biological repeats. **(D)** HDF-tet^on^-GFP-progerin expressing progerin for 8 days (and control) were incubated with calcitriol (100nM) or vehicle and transfected with poly(dA:dT). cGAMP was measured by ELISA. Data represent average±SEM of 3 biological repeats. **(E)** Percentage of cells positive for ^S366^p-STING and percentage of ^S366^p-STING at the PNC in HDFs and **(F)** NF and HGPS fibroblasts. The mean of each of 3 biological repeats is shown. In each experiment, ∼25 image fields were analyzed (n>200). The average±SEM of all fields in 3 experiments is represented. Statistical significance determined by two-way ANOVA.

### Vitamin D signaling regulates the canonical and non-canonical cGAS-STING pathways

We previously demonstrated the robust rejuvenation effects of vitamin D treatment in progeria cells and mice (47-49). The active hormonal form of vitamin D, calcitriol, reduces genomic instability, mitochondrial dysfunction, autophagy defects, reduced proliferation, and sterile inflammation, in progeria cells (47). Here, we determined whether calcitriol has an effect in the activation of the canonical cGAS-STING pathway in progeria and senescent cells (**Supplemental Figure 4D**,**E**). We triggered the canonical pathway with poly(dA:dT). In IR-induced senescent fibroblasts and in late passage fibroblasts, calcitriol improves cGAMP production in response to poly(dA:dT) (**Figure 4C**). Similarly, calcitriol normalizes the production of cGAMP in progerin-expressing cells transfected with poly(dA:dT) (**Figure 4D**). Moreover, calcitriol rescues in part STING behavior in progeria cells. As shown in the images (**Supplemental Figure 4F)** and the graph (**Figure 4E**), calcitriol increases the progeria cells positive for ^S366^p-STING and PNC trafficking upon activation by poly(dA:dT). Calcitriol treatment also normalizes STING behavior in HGPS patients-derived fibroblasts (**Figure 4F and Supplemental Figure 4G**).

These data indicate that vitamin D/calcitriol signaling not only plays a role repressing sterile inflammation via the non-canonical cGAS-STING pathway in progeria; but also rescues the ability of aging/progeria cells to trigger a robust canonical cGAS-STING pathway in response to synthetic dsDNA. Thus, calcitriol regulates both the canonical and non-canonical cGAS-STING pathways, providing a molecular mechanism to tune these pathways in response to infection or disease.

### Targeting the non-canonical STING pathway ameliorates organismal aging

Given the improvement of cellular aging phenotypes by H151, we determined how STING inhibition impacts health and lifespan of progeria mice. We used *Lmna*^*G609G/G609G*^ mice (50), which recapitulate human tissue degeneration phenotypes, especially the loss of VSMCs in the aorta and of WAT (lipodystrophy). H151 treatment from postnatal day 50 (**Figure 5A**) slowed-down the loss of body weight (**Figure 5B**), and significantly increased the lifespan of progeria mice. A median survival of 152 days in mice treated with H151 represents a 26% increase when compared to 120.5 days in untreated progeria mice (**Figure 5C**). Male and female mice treated with H151 experienced similar improvements in body weight (**Supplemental Figure 5A**,**C**) and lifespan (**Supplemental Figure 5B**,**D**).

**Figure 5.**
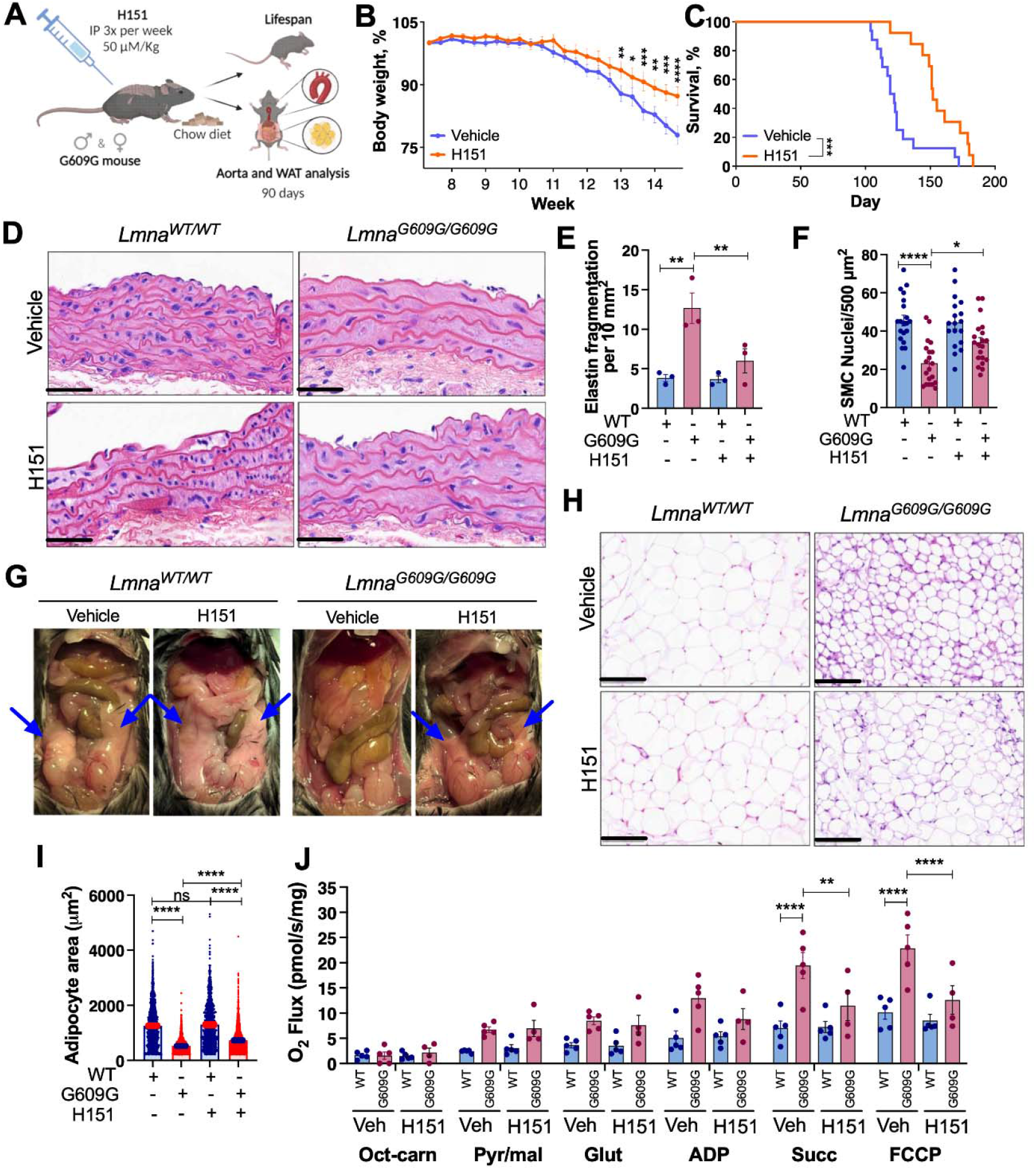
Pharmacological inhibition of STING improves health and lifespan of progeria mice. **(A)** Schematic of the treatment of *Lmna*^*G609G/G609G*^ (G609G) and *Lmna*^*+/+*^ (WT; wild-type) with H151 used for lifespan studies or euthanized at 90 days of age for tissue analysis. Female and male mice were treated, and aorta and WAT were analyzed. **(B)** Body weight of mixed male and female G609G mice treated with H151 (IP 50 μM/Kg, n=13) or Vehicle (n=16) and fed a standard chow diet. Body weight was monitored three times per week. **(C)** Kaplan–Meier survival curves of mixed male and female G609G mice fed chow diet and treated with H151 (n=13) or vehicle (n=16). Median survival of chow-fed G609G mice is 120.5 days for vehicle-treated mice and 152 days for H151-treated mice (26% improvement). **(D)** H&E staining of aortic arch from chow fed G609G mice treated with H151 or vehicle (90 days of age). Scale bars: 50 μm **(E)** Quantification of elastin fragmentation frequency per 10 mm2 (n=3 mouse aortas per group). **(F)** Quantification of smooth muscle cell (SMC) nuclei per 500 μm2 in six different areas of each independent mouse (n=3 mouse aortas per group). **(G)** Pictures show WT and G609G mice treated with vehicle or H151 and fed a standard chow diet (90 days of age). Blue arrows highlight the presence of epidydimal fat in each mouse. **(H)** Histology of epidydimal WAT stained with H&E (n=3 mouse eWAT per group). Scale bars: 100 μm. **(I)** Quantification of adipocyte size in μm2 of approximately 500 adipocytes per sample (n=3 mice). **(J)** Mitochondrial respiration assessment in eWAT from G609G and WT mice treated with vehicle or H151 using Oroboros instrument. Graphs show average ± SEM of oxygen flux after additions of octanoyl-l-carnitine (Oct-carn); pyruvate and malate (Pyr/Mal); glutamate (Glut); adenosine diphosphate (ADP), succinate (Succ), and FCCP (n = 4-5).

Next, we monitored the impact of STING inhibition in tissue degeneration. Treatment with H151 was initiated at postnatal day 50, and mice were euthanized, and tissue collected at day 90. Aging phenotypes were evident in the vasculature and WAT, resembling HGPS patients’ phenotypes. As shown in the histology of aortas (**Figure 5D**), progeria mice exhibit increased elastin fragmentation (**Figure 5E**), and reduced number of VSMCs (**Figure 5F**), which are improved by H151 treatment. Progeria mice exhibit severe lipodystrophy (loss of visceral and subcutaneous WAT) (**Figure 5G**). The beneficial impact of H151 on WAT was evident at necropsy, where we noted the persistence of visceral fat in the H151-treated progeria mice vs untreated mice. Moreover, WAT histology (**Figure 5H**) revealed reduced adipocyte area (size) (**Figure 5I**) that was improved by H151. We previously reported that WAT of progeria mice has increased mitochondrial activity/respiration compared to wild-type, consistent with “browning” (42). We show that H151 treatment normalizes mitochondrial function in WAT of progeria mice (**Figure 5J**). In contrast to the hyperactivity of mitochondria observed in WAT, we find reduced mitochondrial respiration in heart tissue (**Supplemental Figure 5E**), and H151 does not improve mitochondrial dysfunction in this tissue. Future studies need to determine the mechanisms whereby STING impacts mitochondrial function and metabolic pathways in different tissues. Our data here indicate that a non-canonical STING pathway drives tissue degeneration in progeria and provides a target for therapeutic intervention to ameliorate some aging-related phenotypes.

## Discussion

The critical role that cGAS-STING pathway plays in inflammation during aging and senescence is indisputable (2, 6, 7, 28, 29, 44). Targeting cGAS or STING reduces inflammation and improves other cellular and organismal aging-associated phenotypes (13, 51). However, the molecular mechanisms involved remain poorly understood. Understanding the cGAS-STING signaling axis in-depth will allow developing selective inhibitors and agonists that manipulate this pathway towards specific therapeutic goals. The current understanding is that genomic instability and mitochondrial dysfunction during senescence/aging lead to buildup of self-DNA in the cytoplasm, which is recognized by cGAS leading to cGAMP production (9). cGAMP binds to STING (10), inducing oligomerization and trafficking from ER to PNC, where STING forms a ternary complex with TBK1 and IRF3, resulting in phosphorylation of the three proteins and activation of IRF3 and NFκB transcriptional programs. This canonical pathway has been well-described in the context of microbial infection. However, whether all these markers of the canonical pathway are conserved in the context of self-DNA accumulation in the cytoplasm is not known. Studies have clearly shown that depletion/inhibition of cGAS or STING reduces inflammation in different types of senescence (2, 28, 29, 33, 44). It is less clear however, whether the enzymatic activity of cGAS or other functions of cGAS are required for driving senescence. It is also not clear whether STING localizes to the PNC and forms the ternary complex with TBK1 and IRF3, resulting in its phosphorylation (28). In fact, recent studies question the activation of the cGAS-STING pathway in response to DNA damage and micronuclei (52, 53) Here, we show that a cGAS- and STING-dependent inflammatory response is activated in aging, senescent, and progeria cells. However, this response is not triggered through the canonical cGAS-STING pathway, as shown by the lack of detectable increase in cGAMP, translocation of STING to PNC, or phosphorylation of STING, TBK1, and IRF3 ternary complex. Instead, a non-canonical cGAS-STING pathway is activated in these cells, characterized by increased presence of STING in the nucleus (NE and chromatin), that plays a major role in cellular and organismal aging in progeria.

Our study provides the new paradigm that STING can induce sterile inflammation in aging without a significant increase in cGAMP production by cGAS. The mechanism/s behind the reduced cGAMP production in senescence/aging/progeria despite the accumulation of cytosolic DNA is not known. cGAS is localized to both, cytosol and nucleus (54, 55). The cytoplasmic pool is considered responsible for responding to foreign/self-DNA in the cytosol and synthesizing cGAMP, while the nuclear pool is kept enzymatically inactive by binding to histones in the nucleosome (50). Histone-bound self-DNA has also been shown to be inhibitory to cGAS activation in cells presenting with micronuclei (MN) generated by genotoxic insults (52). This is due to the acidic patch of H2A and H2B outcompeting dsDNA for binding cGAS (56). Whether the accumulation of MN in progeria and senescent cells sequesters cGAS and inhibits its activity is not known. However, a clear difference between the cells that exhibit MN due to genotoxic insults and progeria/senescent cells, is the lack of activation of the IFN response in the former, and the robust activation in aging cells. Thus, the mechanism of inhibition of cGAS might be different in both contexts. Interestingly, a study performed in human monocyte-derived dendritic cells revealed that nuclear-localized cGAS can synthesize low levels of cGAMP (200-fold lower than following transfection with dsDNA) and induce innate immune activation (57). Further studies are needed to determine whether cGAS localization and functions are altered during the aging process, and whether these are linked to the changes in cGAMP production and STING behavior identified here.

It is well-established that STING is activated by binding to cGAMP produced by cGAS. A recent study in cortex of aged mice shows that the increase in cytoplasmic DNA, attributed to DNA damage, leads to an increase in cGAS levels and cGAMP production. However STING:TBK1:IRF3 complex is not activated in this tissue (58). These data suggest an uncoupling of cGAS activity from STING activation. Thus, cGAS and STING could also play independent roles in aging, which remains to be explored. For instance, the DNA sensor protein IFI16 has been shown to activate STING in a cGAS/cGAMP-independent manner in etoposide-treated cells (18). Upon DNA damage (etoposide), ATM and PARP-1 facilitate the formation of an alternative STING complex consisting of IFI16/p53/TRAF6/STING, which causes ubiquitylation of STING and activation of the NFκB pathway without exit from the ER. This non-canonical STING pathway triggers an innate immune gene expression profile that differs from that induced by the canonical cGAS-STING pathway (18). However, another study showed that IFI16 can cooperate with cGAS to activate STING during DNA sensing in human keratinocytes, and that IFI16 promotes STING phosphorylation and translocation (17). Thus, even though the current dogma states that STING signaling depends on its trafficking to PNC, these studies and our data indicate that STING can signal and induce inflammation from other cellular compartments.

It will be important to define whether the altered behavior of STING in aging/senescent/progeria cells is accompanied by new partnerships in the nucleus and in chromatin, which might reveal new ways whereby STING regulates the inflammatory/IFN response and potential novel functions of STING in the nucleus. It was recently reported that nuclear STING participates in transcription; a function that competes with signaling from the canonical cGAS-STING pathway (59). Other reports have linked nuclear STING with genomic stability/instability. Nuclear STING protects breast cancer cells from DNA instability via a non-canonical, cGAS-independent pathway that promotes DNA repair and resistance to genotoxic agents (60). In this context, STING is partially localized at the NE, and interacts with proteins in the DNA damage response, including the DNA-PK complex. STING also modulates nascent DNA metabolism at stalled replication forks, with STING activation causing replication stress and nascent DNA degradation (61). Another study shows that STING protects replication forks from a distance, by triggering a Ca^2+^-regulated pathway at the ER that prevents nascent DNA degradation (62). Thus, the effects of STING on the regulation of genome function and maintenance are likely to be cell- and context/insult-specific, and their in-depth characterization is necessary to fully understand STING’s multifaceted functions.

The data presented here targeting STING *in vivo* with inhibitor H151 demonstrates a significant improvement of tissue homeostasis in progeria mice, reducing the dramatic loss of VSMCs in the aorta and inducing persistence of WAT. This is in line with a study showing that STING promotes low-grade inflammation and functional decline in aged mice (>24 months) (57). Inhibition of STING with H151 reduces inflammation in peripheral organs and brain, with significant improvement in learning and memory performance. In the brains, the authors found STING and TKB1 phosphorylation, and higher cGAMP production than in young brains, primarily in microglia. This indicates that in the brain/microglia of old mice there is activation of the canonical cGAS-STING pathway. Interestingly, production of cGAMP was shown in mouse tail fibroblasts expressing constitutively active cGAS (does not bind to nucleosomes), leading to a robust activation of ISGs. Altogether, these data suggest that cGAMP production might differ among cell types and be regulated by cGAS binding to nucleosomes. Further studies are needed to determine whether the reduced ability to activate the canonical cGAS-STING pathway in human fibroblasts aging in culture is conserved in other cell types and the role that chromatin binding plays in this phenotype.

Our finding that the ability to activate the canonical cGAS-STING pathway is diminished as cells “age” in culture and enter senescence has important implications for therapy development. The cGAS-STING pathway is associated with a growing list of aging-related diseases; considered to be pathogenic in some cases (autoimmune, neurodegenerative, and cardiovascular), and beneficial in others (response to viral infections and cancer treatments) (63). Unveiling the mechanisms responsible for the changes in STING behavior during aging, signaling preferentially through the non-canonical pathway, could provide new therapeutic targets to delay cellular senescence and improve tissue function, to ameliorate tissue degeneration, to slow-down cancer progression, to respond to viral infection, or to treat IFN-driven diseases. This study shows that one of the mechanisms regulating the cGAS-STING pathway in aging is the vitamin D (calcitriol) receptor (VDR) signaling pathway. Calcitriol treatment of aging/progeria cells increases the production of cGAMP and the ability of phosphorylated STING to properly localize to the PNC in response to synthetic DNA. Based on our previous findings that VDR levels decrease during aging and in progeria (47), we propose that VDR deficiencies with age contribute to the alterations in STING behavior and the decreased activity of cGAS. Calcitriol treatment, which increases VDR levels (47), rescues the ability of progeria/aging cells to activate the canonical cGAS-STING pathway in response to synthetic DNA.

## Materials and Methods

### Cell culture

Human dermal fibroblasts (de-identified) containing doxycycline-inducible GFP-progerin construct were cultured at 37°C in DMEM supplemented with 10% FBS and 1% antibiotics/antimycotics and treated for 4-8 days with vehicle (DMSO) or 0.5-1µg/mL doxycycline. Cells were treated 2 days with 0.5µM H151 prior to doxycycline addition. Cells were subcultured when 90% confluent and H151-supplemented media was changed every 2 days. For cGAS inhibition, cells were treated with G140 (5 or 10 µM) for 6 days of progerin induction. The medium was replenished every 24h. Skin fibroblasts from HGPS patients with classic mutation and from their parents (normal fibroblasts, NF) were obtained from Progeria Research Foundation. The HGPS mutant HGADFN164 is from a 4-years-old patient, *LMNA* exon 11 heterozygous c.1824C>T (p.Gly608Gly). NF are from a 53-year-old parent of an HGPS patient, negative for the mutation in the *LMNA* Exon 11. These cells were cultured under 5% CO_2_ and 3% or 20% O_2_ at 37°C in DMEM, 15% FBS, and 1% antibiotics/antimycotics. For inducing senescence, cells were irradiated with 20 Gy and allowed to grow for 3 weeks before being collected.

**Immunofluorescence, Quantitative Reverse-Transcription PCR, Western blot, Calcitriol treatment, and High-resolution respirometry and Histology**. Described in Supplemental Information.

### cGAMP assessment

For cGAMP measurement, cells transfected with 1μg/ml of poly (dA:dT) for 4 hr were lysed in RIPA buffer followed by centrifugation at 14000 x g at 4°C for 15 min. The supernatant was used for cGAMP ELISA assay according to manufacturer’s instructions (2’3’-Cyclic GAMP ELISA Kit, ThemoFisher). cGAMP concentration is shown for 100 μg of protein.

### Mice

Animal experiments were performed in conformity use of laboratory animals and approved by the IACUC at Saint Louis University (protocol #2299). HGPS mouse models (*Lmna*^*G609G/G609G*^) were generated in the laboratory of Carlos Lopez-Otin. The mice used in this study were housed in a pathogen-free facility at 23°C with food and water provided *ad libitum* under a 12:12 light-dark cycle. All animals were fed a regular chow diet and littermates of the same sex were randomly assigned to experimental groups.

Female and male *Lmna*^*G609G/G609G*^ (G609G) mice were treated with 0.5 μM H151 or vehicle (5% Tween 80 in PBS) three times a week through intraperitoneal injection, starting at day 50 of age and continued up to the humane endpoint. Chow-fed G609G mice receiving vehicle or H151 were sacrificed at day 90 of age and tissues were collected for functional assays and histological analysis.

### Statistical analysis

One-way or two-way ANOVA was used for the experiments in which more than two comparisons were performed; if p < 0.05, significance between groups was determined using post hoc analysis. For comparisons between groups, Tukey HSD test was used. Error bars displayed throughout the manuscript represent SEM and were obtained from replicates of each biological sample unless otherwise indicated. Survival curves were generated using the Kaplan–Meier method and log-rank test to determine statistical significance between survival differences. GraphPad Prism 8.02 was used for the calculations. In all cases, differences were considered statistically significant when p < 0.05 (*p < 0.05, **p < 0.01, ***p < 0.001, ****p < 0.0001).

## Acknowledgments

Funding for this work from NIH grants RAG082759A, RAG076145A, and R01AG058714. L.S. funded by Glenn Foundation for Medical Research Postdoctoral Fellowship in Aging Research (PD24164). R.C.F. and B.T.C. supported by the Doisy Fund of the Edward A. Doisy Department of Biochemistry and Molecular Biology.

## Author Contributions

Rafael Cancado de Faria, Lilian Silva, and Elena Shashkova performed all the experiments. Barbara Teodoro-Castro contributed to early stages of the project. Kyle McCommis provided guidance monitoring mitochondrial function. Susana Gonzalo supervised the work.

## Competing Interest Statement

No competing interests.

## Notes

### Competing Interest Statement

The authors have declared no competing interest.

