## Supplemental Methods and Figures for "A non-canonical cGAS-STING pathway drives cellular and organismal aging"

### Supplemental Information

#### Supplemental Materials and Methods

##### Immunofluorescence

For immunofluorescence, cells were plated onto coverslips and allowed to fully attach. After treatments with calcitriol or poly(dA:dT), cells were fixed either with 4% paraformaldehyde (Ki67 staining) for 20 min at R.T. or methanol (STING, p-STING and p65 staining) for 10 min at R.T. Cells were then washed three times with PBS before permeabilization with 0.5% Triton X-100 for 20 min at R.T. Following permeabilization, coverslips were washed 3 times with PBS and blocked 1 h at 37 °C with 2% BSA/PBS. Cells were then incubated with primary antibody anti-Ki67, p65, STING or p-STING in 1% BSA/PBS overnight at 4°C. After incubation, coverslips were washed 3 times in PBS and incubated with secondary anti-rabbit Alexa Fluor 488 (1:1000) or anti-rabbit Alexa Fluor 594 (1:1000) in 1% BSA/PBS for 2 hours in a humid chamber at R.T. Cells were washed 3 times with PBS and counterstained with DAPI. Slides were allowed to dry for 2 hr at R.T. in the dark and stored overnight at 4°C before visualization with a Leica DM5000B microscope. Images were acquired using 63× oil objective lenses with a Leica DFC350FX digital camera and the Leica Application Suite. On average, fifteen-thirty images per sample were obtained and, at least, 200 cells per genotype were quantitated for the analysis of STING/p-STING and p65 localization or Ki67 positivity.

##### Quantitative Reverse-Transcription PCR

Cellular RNA was purified using Direct-zol RNA MiniPrep kit according to manufacturer instructions (Zymo Research). Following RNA extraction, 0.5 µg total purified RNA was used for reverse transcription reactions to obtain cDNA using SuperScript VILO cDNA Synthesis kit. For quantitative PCR, Power SYBR Green PCR Master Mix was used with the 7500HT Fast Real-Time PCR system. Reactions were performed in triplicate and ACTB and GAPDH were used as controls. Cycle thresholds from target genes were compared to determine relative quantitative measurements. Oligo sequences and target genes are listed in the Supplemental materials.

##### Western blot

Cells were lysed with RIPA buffer (50 mM Tris pH 7.5, 1% NP40, 0.25% deoxycholic acid, 1 mM EDTA) for 30 min under constant rotation at 4°C. Cell lysates were centrifuged at 14,000 g for 15 min at 4°C to remove cell pellet. Following centrifugation, 50-60 µg of total protein in the supernatants were mixed with Laemmli loading buffer and incubated at 95°C for 5 min prior electrophoresis. Equal protein amounts were separated by SDS-PAGE on a 4-15% Criterion TGX Gel (Bio-Rad) followed by transferring to a nitrocellulose membrane using the Trans-Blot Turbo system (Bio-Rad). After transferring, all membranes were blocked with 5% BSA or 5% milk in 0.1% Tween 20/TBS for 1 hour at R.T., and incubated with primary antibody diluted in blocking solution overnight at 4°C. Following incubation, membranes were washed three times with 0.1% Tween 20/TBS and incubated with secondary antibodies (anti-rabbit, anti-mouse, or anti-chicken, HRP conjugate) for 2 h at R.T. Membranes were washed three times with 0.1% Tween 20/TBS and developed using Pierce ECL Western Blotting Substrate (Thermo Scientific). Western blot images were acquired using SYNGENE PXi instrument and protein bands densitometry was quantified using ImageJ. Densitometry values were normalized by one of the housekeeping loading controls. In case of phosphorylated proteins, quantification of phosphorylation was obtained over unphosphorylated protein levels.

To monitor cGAS levels cells were lysed with RIPA buffer above containing 0.5% SDS, for 30 min under constant rotation at 4°C. Then, cell lysates were sonicated for 5 min and centrifuged at 14,000 g for 15 min at 4°C to remove cell pellet.

**Calcitriol treatment**

Cells were supplemented with 100 nM calcitriol (1α,25-dihydroxyvitamin $D_3$) where indicated. Aliquots of calcitriol were resuspended in FBS and diluted in DMEM to a final concentration of 10% FBS for HDF-*tet*$^{on}$-GFP-progerin or 15% FBS for NF and HGPS fibroblasts.

For replicative senescence experiments, early-passage (passage 22) and late-passage (passage 35) fibroblasts were treated with 100 nM calcitriol for seven days before stimulation of the cGAS-STING pathway with 1 µg/mL poly(dA:dT) for four hours. For DNA damage-induced senescence, early-passage fibroblasts (passage 22) were exposed to 20 Gy ionizing radiation and subsequently cultured for 14 days. Irradiated cells were cultured with 100 nM calcitriol for the following seven days and, on day 21, stimulated with 1 µg/mL poly(dA:dT) for four hours. For HDF-*tet*$^{on}$-GFP-progerin, the cells were pre-treated with calcitriol for two days, followed by progerin induction using 1 µg/mL doxycycline alongside continued treatment with 100 nM calcitriol for an additional eight days. In experiments using fibroblasts derived from HGPS patients (passage 12), the cells were treated with 100 nM calcitriol for seven days. Prior to cell collection or fixation, the cGAS-STING pathway was stimulated with 1 µg/mL poly(dA:dT). In all experiments using calcitriol, the media was changed every 2–3 days, and cells were passaged upon reaching 90% confluence. Cells were harvested or fixed after synthetic DNA stimulation for cGAMP assessment via ELISA or $^{S366}$p-STING immunofluorescence, respectively.

##### High-resolution respirometry

High-resolution respirometry experiments using an Oroboros Oxygraph O2k FluoRespirometer (Oroboros Instruments) were performed as previously reported (41). Briefly, after excision, the rostral zone of eWAT samples were immersed in BIOPS buffer (50 mM MES, 10 mM EGTA, 6.56 mM $MgCl_2$, 0.5 mM DTT, 20 mM imidazole, 5.77 mM ATP, and 15 mM phosphocreatine, pH 7.1). Approximately 1 mg WAT was then placed in prewarmed Oroboros chambers with mitochondrial respiration solution MiR05 (3 mM $MgCl_2$, 0.5 mM EGTA, 20 mM taurine, 60 mM k-lactobionate, 10 mM $KH_2PO_4$, 110 mM sucrose, 20 mM HEPES, and 1 g $L^{-1}$ BSA, pH 7.1). Oxygen was provided to ensure $O_2$ availability within the eWAT permeabilized with 2 µM digitonin. For $O_2$ flux measurement, different substrates were sequentially added: 1.5 mM octanoyl carnitine (OC), 5 mM pyruvate, 10 mM glutamate and 2 mM malate, 20 mM ADP, and 20 mM succinate, followed by 3 pulses of 0.5 µM FCCP. $O_2$ flux per mass was recorded using DatLab 7.4 software.

For mitochondrial respiration analyzes in heart tissue, left ventricle was excised from the mice and immersed in an ice cold solubilization solution containing 5 mg $mL^{-1}$ saponin in BIOPS buffer for 20 minutes at 4 °C. Samples were then immersed in the MiRO5 buffer for an additional 20 minutes. 1-mg pieces of solubilized heart were placed into the Oroboros chambers filled with MiRO5 buffer, 3 mg mL-1 creatine and 2 uM blebbistatin. Chambers were oxygenized and $O_2$ flux measurements were performed with substrate additions as previously described for eWAT.

##### Histology

Histological analysis was performed as in (41). Mice were euthanized 90 days of age and aortic arch and epididymal WAT were collected, washed with PBS and incubated in 10% formalin. Tissue processing and staining with hematoxylin and eosin were performed at HistoWiz. The aortic arch was collected, cleaned from fat tissue, and fixed in 10% formalin. The number of smooth muscle cells at the aortic arch was obtained by counting the number of nuclei per 500 $\mu m^2$ in the media layer. For histological analysis of eWAT, adipocyte area was calculated using ImageJ.

#### Supplemental Figures and Tables

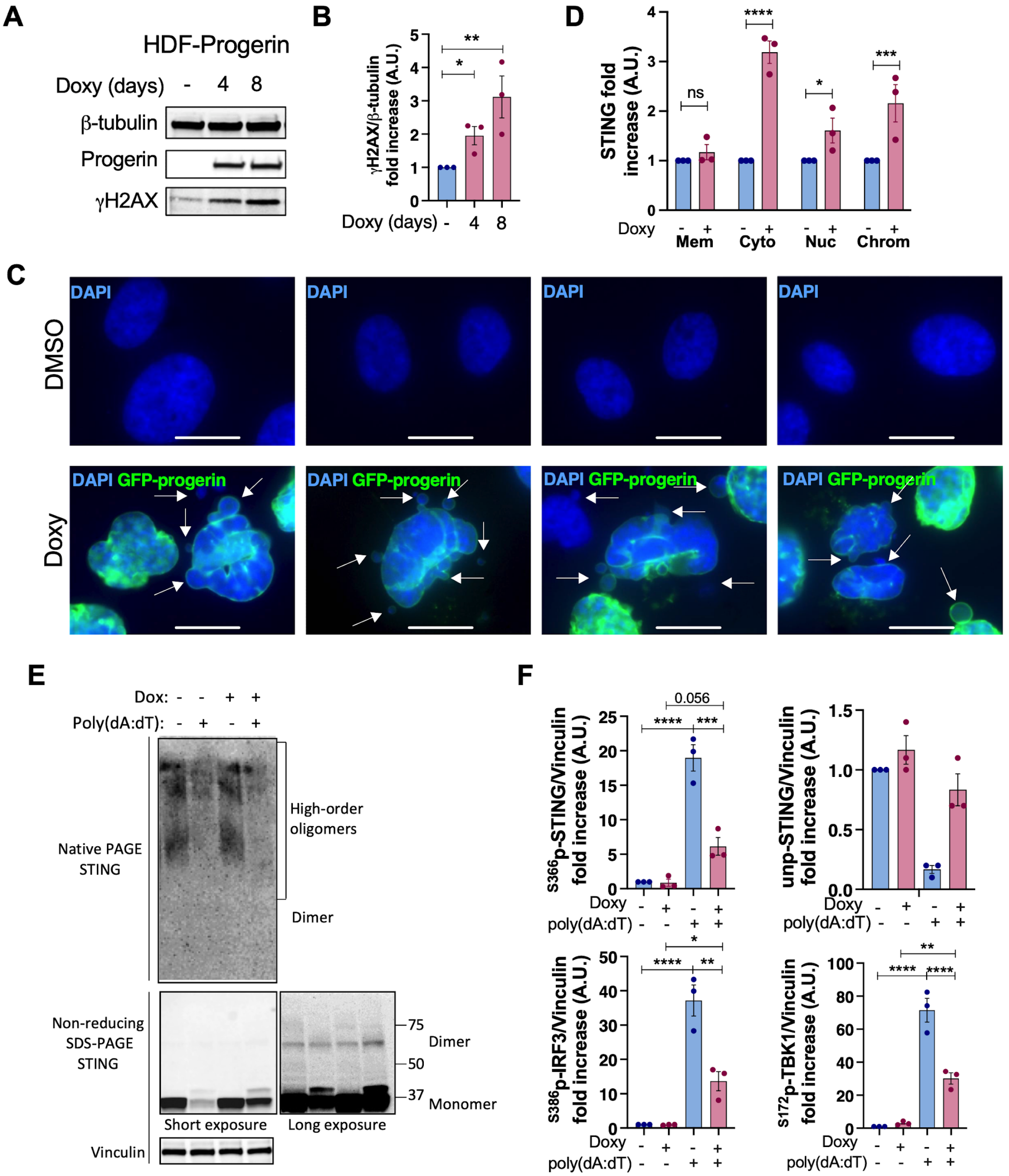


**Supplemental Figure 1. Accumulation of cytosolic DNA, DNA damage, and altered STING behavior in progeria cells.** (**A**) Doxycline-induced progerin expression in HDFs leads to increased levels of DNA damage, monitored by γH2AX immunoblotting. (**B**) Densitometry of γH2AX levels upon 4-8 days of induced expression of progerin. Average±SEM of 3 independent experiments. (**C**) Images of cells expressing GFP-progerin show nuclear envelope deformation/protrusions, chromatin fragments in the cytoplasm and micronuclei. Scale bars: 10 μm. (**D**) Densitometry of STING signal in 3 independent subcellular fractionation experiments, normalized to the corresponding marker of each fraction. Note the increase in STING bound to chromatin in progerin-expressing cells. (**E**) Control and progerin-expressing HDF (+Dox) were treated with vehicle or poly(dA:dT), and whole cell protein lysates were collected and analyzed in different PAGE conditions. Top immunoblot shows STING signal in native PAGE. Note the different oligomers in vehicle treated cells, and the presence of STING at high-order oligomers in poly(dA:dT)-treated cells. No differences are observed between control and progerin-expressing cells in terms of oligomers formation. Bottom immunoblots show monomeric STING in non-reducing SDS-PAGE. Left panel shows how poly(dA:dT) triggers the phosphorylation (upper STING band) and degradation of STING in control cells. In progerin-expressing cells, there is less degradation and accumulation of unphosphorylated STING. Right panel is a higher exposure showing that there are no differences in dimer formation between the different conditions. Vinculin was used as loading control. (**F**) Densitometry of $^{S366}$p-STING, $^{S172}$p-TBK1, $^{S386}$p-IRF3, and unphosphorylated STING in control and progerin-expressing HDF that are treated +/- poly(dA:dT). Average±SEM of 3 biological repeats. Statistical analysis by one-way ANOVA.

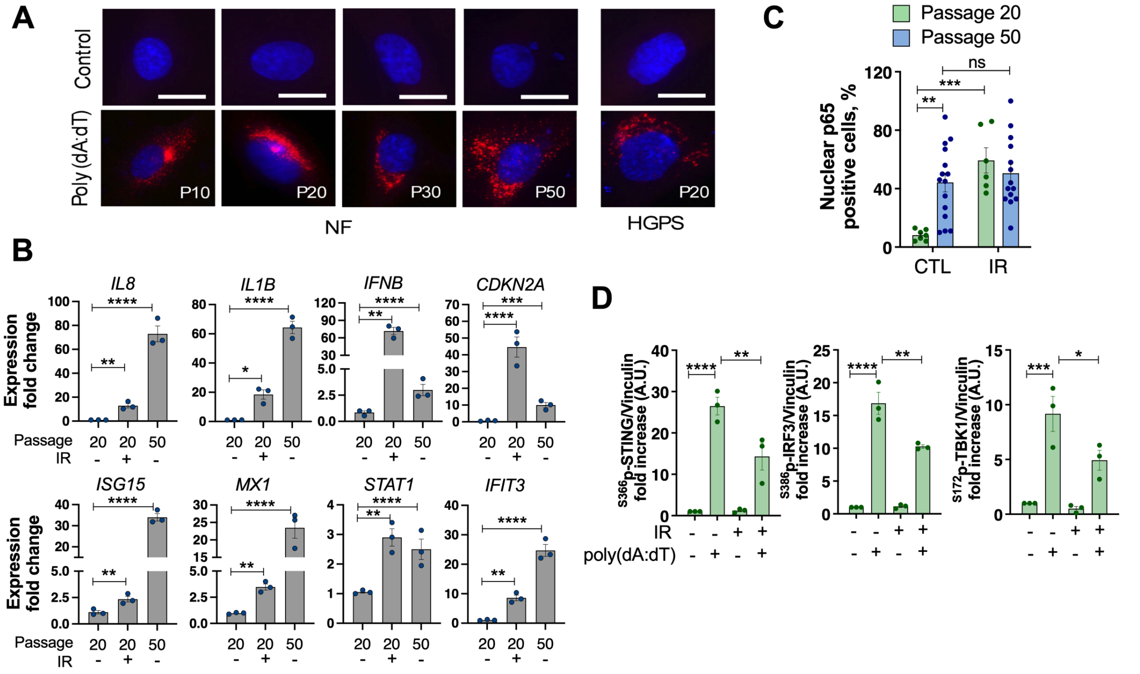


**Supplemental Figure 2. Replicative and DNA damage-induced senescent cells show reduced response to synthetic DNA.** (**A**) Related to Figure 2D. IF images of $^{S366}$p-STING and DAPI showing NF at passage 10, 20, 30, and 50, and HGPS at passage 20 that have been transfected with poly(dA:dT) to activate the canonical cGAS-STING pathway. Note the absence of $^{S366}$p-STING label in cells that are treated with vehicle, and the reduced localization at PNC as NF are passaged in culture, mirroring HGPS fibroblasts. Scale bars: 10µm. (**B**) qRT-PCR of transcripts encoding inflammatory cytokines (*IL8*, *IL1B,* and *IFNB*), ISGs (*ISG15, MX1, STAT1,* and *IFIT3)*, and p16 (*CDKN2A*), from NF passage 20 and 50, and NF passage 20 irradiated to undergo senescence. (**C**) IF with p65 antibody in early passage (P20), late passage (P50, replicative senescence), and IR-induced senescent cells. Percentage of cells with nuclear translocation of p65, indicative of activation of NFκB pathway, was quantitated. In every condition, ~15 image fields (>150 cells) were analyzed. The average±SEM of all fields is shown. (**D**) NF control and senescent (3 weeks post-IR) were treated with poly(dA:dT) for 4 hrs. and $^{S366}$p-STING, $^{S172}$p-TBK1, and $^{S386}$p-IRF3 were monitored by immunoblotting (Figure 3G). Densitometry from 3 biological repeats (IR-induced senescent cells generated 3 times). Graphs show average± SEM. Statistical analysis by one-way ANOVA. Note the reduced ability of senescent fibroblasts to activate these markers of the canonical cGAS-STING pathway, compared to early passage proliferating fibroblasts.

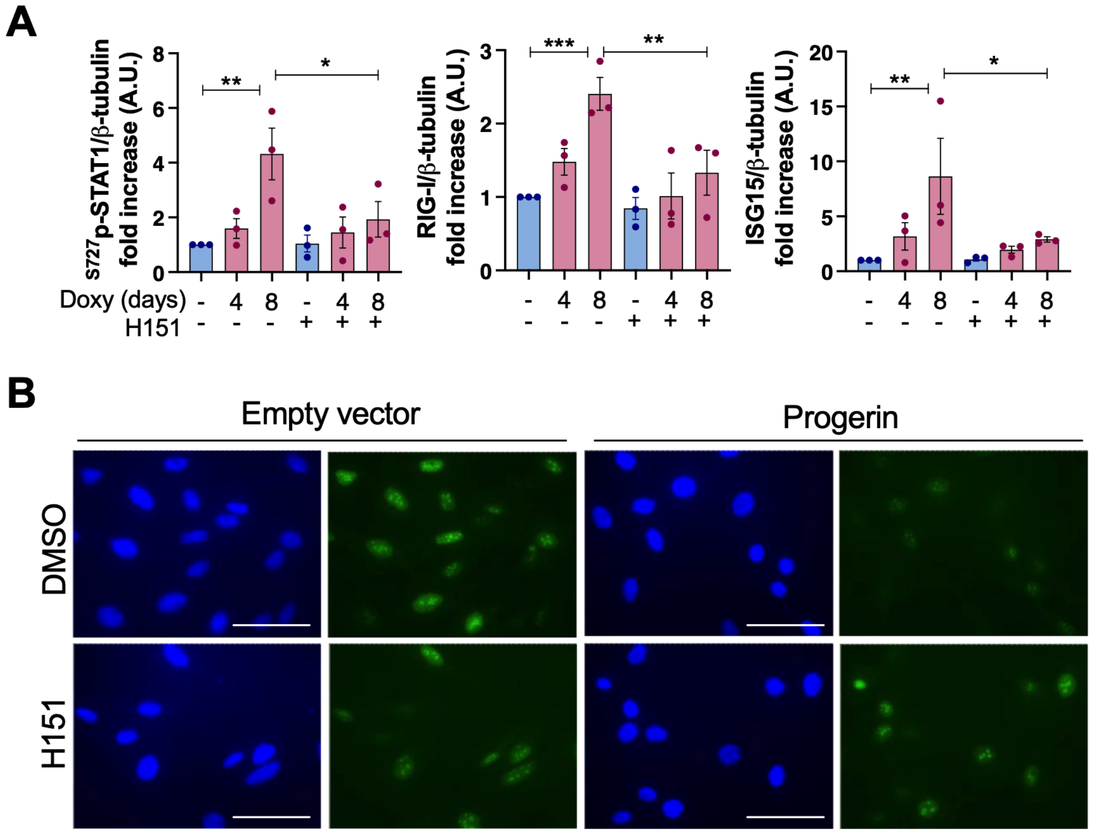


**Supplemental Figure 3. STING inhibition improves proliferation and reduces inflammation. (A)** HDF-teton-GFP-progerin induced to express progerin for 4-8 days were treated with vehicle (DMSO) or H151 (STING inhibitor) during the 4-8 days (0.5μM H151). Densitometry from three independent experiments (average±SEM) showing markers of sterile inflammation/IFN response: p-STAT1, RIG-I and ISG15. Normalized to β-tubulin (loading control). **(B)** IF with Ki67 antibody to monitor proliferating fibroblasts expressing progerin or empty vector control (Ki67 positivity). Scale bars: 50 μm.

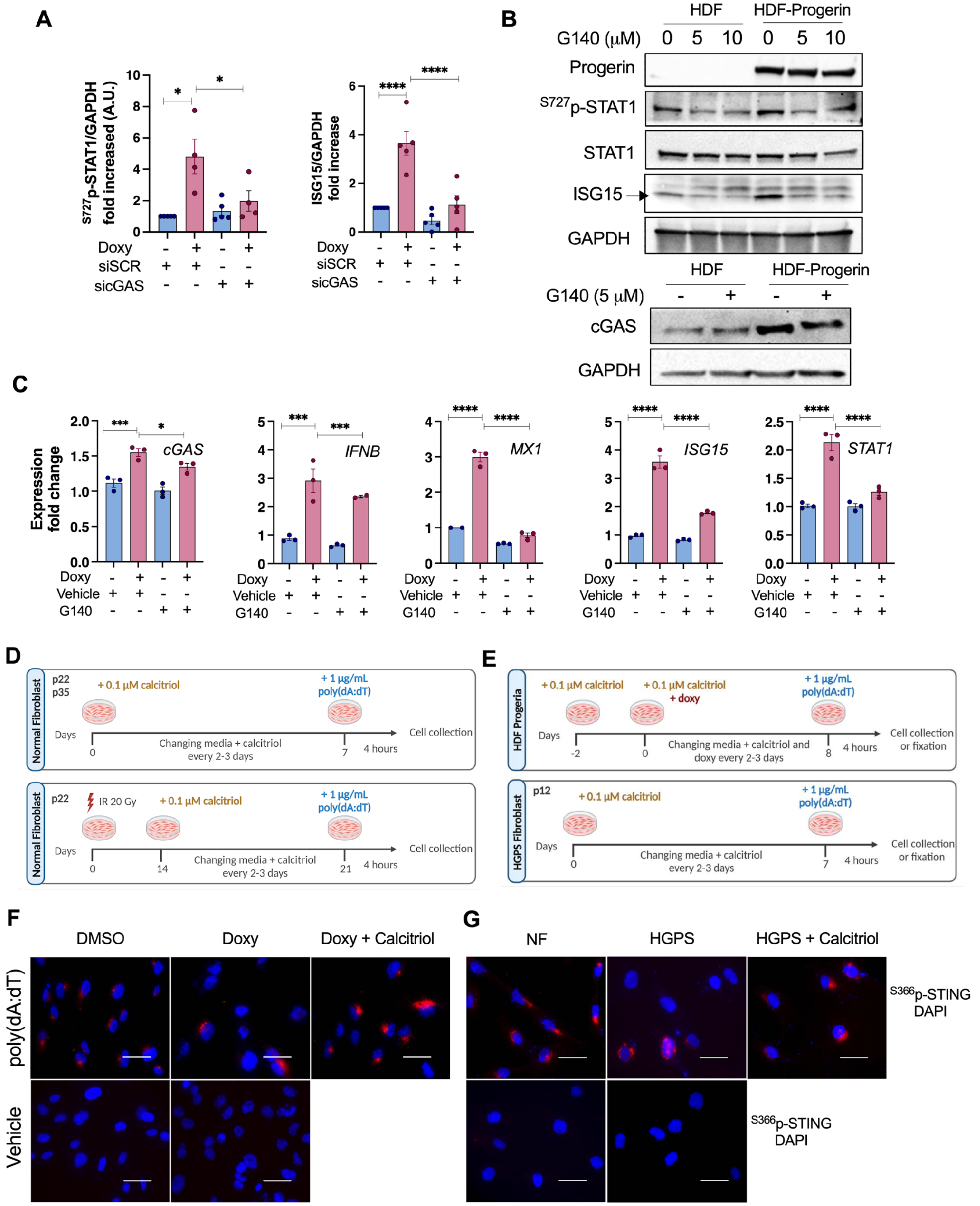

**Supplemental Figure 4. Effect of cGAS inhibition and calcitriol treatment in progerin-expressing cells.** (**A**) Densitometry from Figure 4A calculated from 5 biological repeats, with average±SEM. HDF-*tet*$^{on}$-GFP-progerin induced to express progerin for 8 days (doxy) were transfected with sicGAS or control siRNA. (**B**) HDF-*tet*$^{on}$-GFP-progerin induced to express progerin for 8 days (doxy) were treated with vehicle (DMSO) or G140 (cGAS inhibitor) during the 8 days of progerin induction (5-10μM G140). Representative blots from three independent experiments showing markers of sterile inflammation/IFN response: STAT1, $^{S727}$p-STAT1, and ISG15. GAPDH was used as loading control. cGAS was also monitored by immunoblotting (bottom panels). (**C**) Same cells as in (B) were processed for qRT-PCR to monitor the levels of cGAS and the expression of inflammatory markers (*IFNB, STAT1, ISG15, and MX1*). Note how the increased expression of inflammatory genes in progerin-expressing cells is ameliorated by cGAS inhibition. Graphs show one biological repeat in triplicate, with average±SEM. (**D**) Scheme of treatment of NF of early passage (P22) and late passage (P35) with calcitriol for 7 days prior to transfection with poly(dA:dT) to activate the canonical cGAS-STING pathway. Cells were collected 4 hrs later. Early passage NF were also irradiated with 20 Gy and grown in culture for 21 days to induce senescence. Calcitriol was added for the last 7 days. Senescent cells +/- calcitriol were transfected with poly(dA:dT) and collected at 4 hrs. (**E**) HDF were treated with calcitriol 2 days prior to progerin expression (doxy) and maintained throughout the experiment. At day 8, HDFs-progerin were treated with poly(dA:dT) and cells collected after 4 hrs. In addition, HGPS fibroblasts of early passage (P12) were treated with calcitriol for 7 days and canonical pathway activated with poly(dA:dT). Cells collected for analysis 4 hrs later. (**F**) HDF-$tet^{on}$-GFP-progerin expressing progerin for 8 days (and control) were transfected with poly(dA:dT). Immunofluorescence pictures of $^{S366}$p-STING (red) and DAPI (blue) in HDFs and (**G**) NF and HGPS fibroblasts. Scale bars: 50µm.

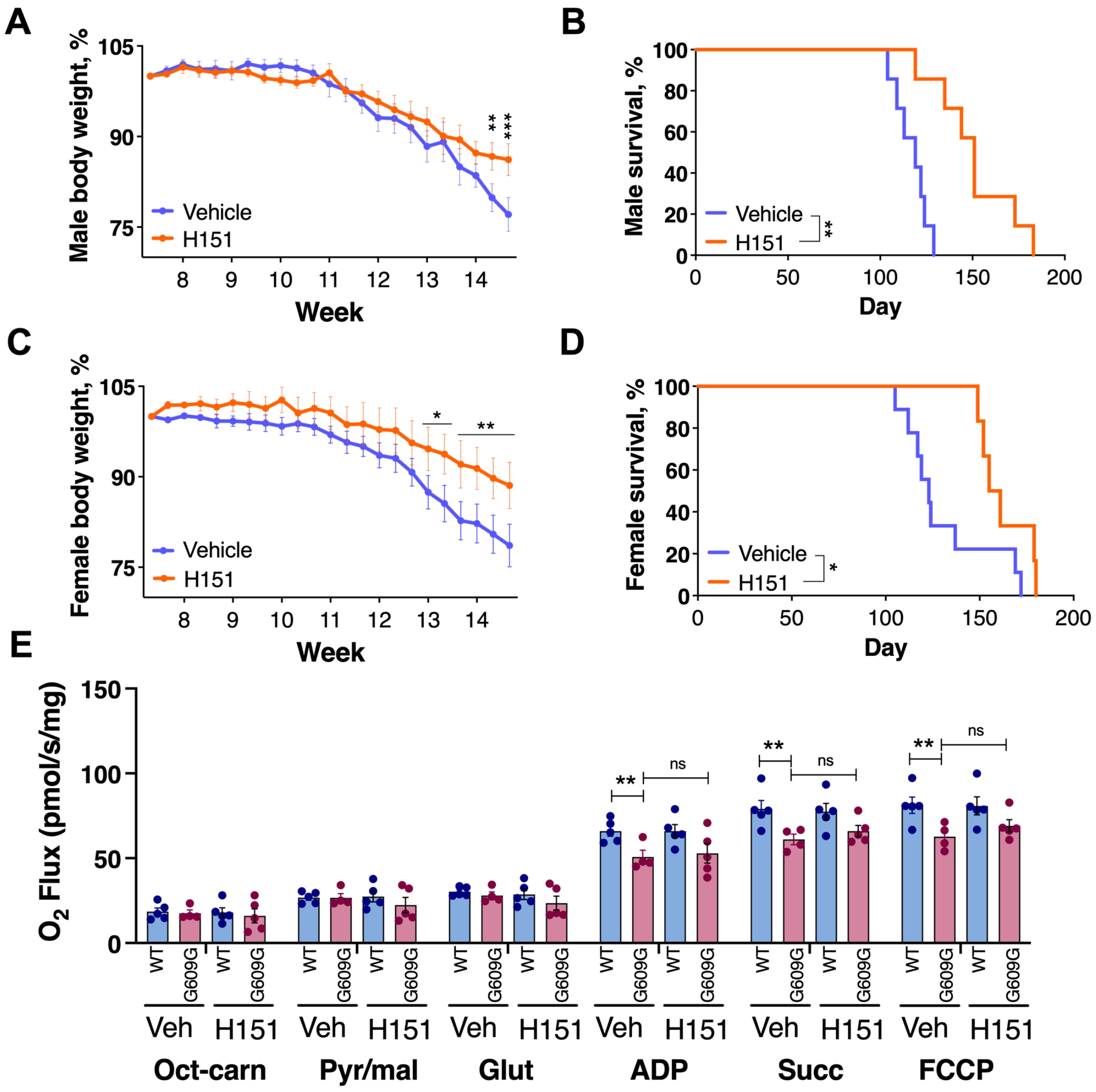


**Supplemental Figure 5. Pharmacological inhibition of STING improves health and lifespan of progeria mice.** (**A**) Male *Lmna*$^{G609G/G609G}$ (G609G) mice were treated with H151 (IP 50 μM/Kg) or Vehicle and body weight was monitored three times per week. (**B**) Kaplan–Meier survival curves of male G609G mice fed chow diet and treated with H151or vehicle. (**C**) Body weight of female G609G mice treated with H151 or vehicle. (**D**) Kaplan–Meier survival curves of female G609G mice fed chow diet and treated with H151or vehicle. (**E**) Mitochondrial respiration assessment in heart tissue from G609G mice treated with vehicle or H151 using Oroboros instrument. Graphs show average±SEM of oxygen flux after additions of octanoyl-l-carnitine (Oct-carn); pyruvate and malate (Pyr/Mal); glutamate (Glut); adenosine diphosphate (ADP), succinate (Succ), and FCCP (n = 4-5).

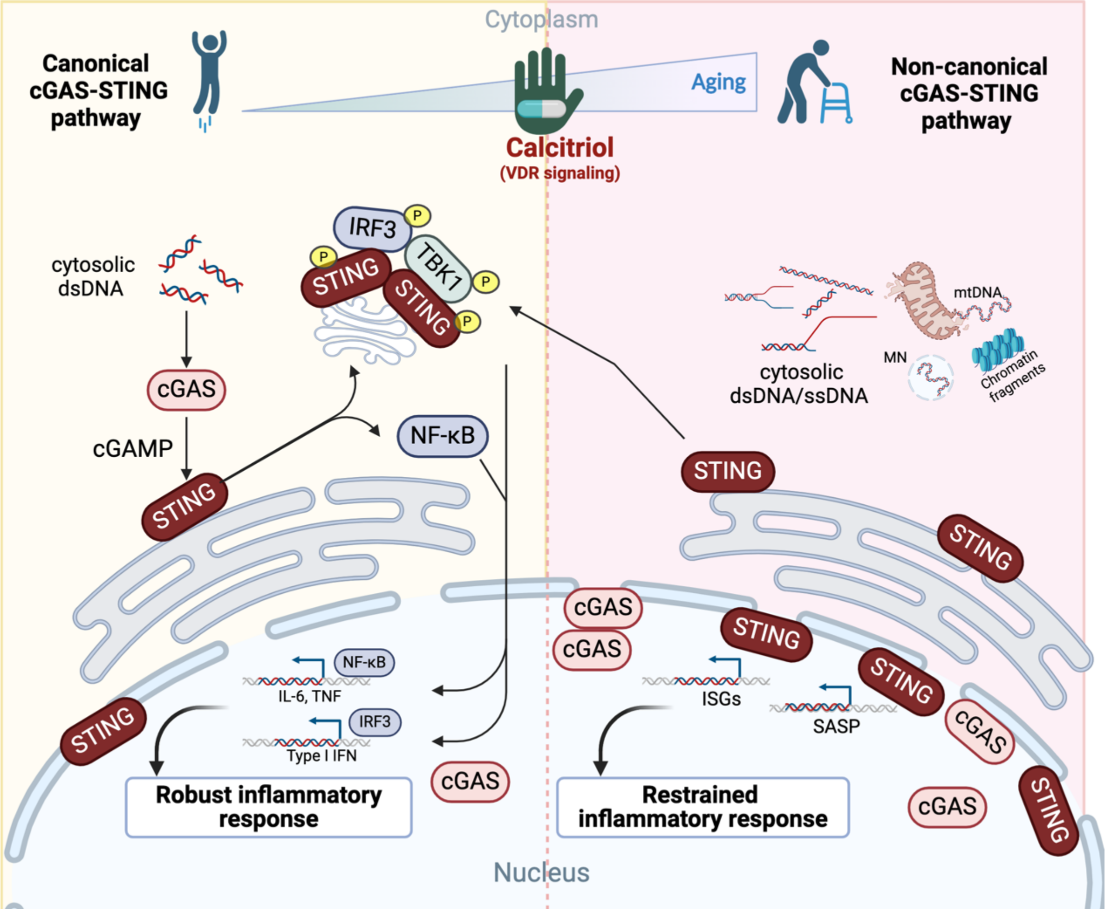


**Supplemental Figure 6. Graphical abstract of activation of a non-canonical cGAS-STING pathway in aging.** Accumulation of cytosolic dsDNA in young cells, such as in the context of microbial infection, activates the canonical cGAS-STING pathway. This pathway is characterized by increased synthesis of cGAMP by cGAS, binding of cGAMP to STING, trafficking of STING to PNC, formation of ternary complex STING/TBK1/IRF3, phosphorylation of STING and other complex proteins, and translocation of IRF3 and NFκB to the nucleus for transcription of pro-inflammatory molecules. During aging, cells progressively reduce their ability to activate the canonical cGAS-STING pathway, despite the accumulation of cytosolic DNA. In aging/progeria cells, cGAS and STING still drive sterile inflammation but via a non-canonical pathway; not involving high cGAMP production, STING phosphorylation or trafficking to PNC. Instead, STING remains associated with ER, NE, and also enriched in chromatin. The non-canonical cGAS-STING pathway plays a major role in cellular and organismal aging phenotypes in HGPS. Importantly, activation of VDR signaling by calcitriol improves the ability of aging/progeria cells to activate the canonical cGAS-STING pathway, while suppressing inflammation through the non-canonical cGAS-STING pathway.

**Supplemental Table 1.** Antibodies and dilutions in this study.

| Western blot | | | |
|---|---|---|---|
| **Antibody** | **Dilution** | **Company** | **Catalog number** |
| β-tubulin | 1-5,000 | Origene | AP31823PU-N |
| Lamin A | 1-3,000 | Abcam | 26300 |
| Progerin | 1-1,000 | Santa Cruz | 81611 |
| H3 | 1-60,000 | Abcam | 1791 |
| ISG15 | 1-1,000 | Santa Cruz | 166755 |
| Ki67 | 1-700 | Cell Signaling | 9129S |
| TBK1 | 1-1,000 | Cell Signaling | 3013S |
| S172 p-TBK1 | 1-1,000 | Cell Signaling | 5483S |
| STAT1 | 1-1,000 | Cell Signaling | 14994S |
| Y701 p-STAT1 | 1-1,000 | Cell Signaling | 7649S |
| S727 p-STAT1 | 1-1,000 | Cell Signaling | 9177S |
| STING | 1-1,000 | Cell Signaling | 13647 |
| S366 p-STING | 1:1,000 | Cell Signaling | 50907 |
| S386 p-IRF3 | 1;1000 | Cell Signaling | 37829 |
| p16 | 1:1,000 | BD Pharmingen | 554079 |
| p65 | 1:1,000 | Cell Signaling | 8242 |
| S536 p-p65 | 1:3,000 | Cell Signaling | 3033 |
| Vinculin | 1:1,000 | Santa Cruz | 73614 |
| Lamin B1 | 1:4,000 | Abcam | 16048 |
| RIG-I | 1-1,000 | Cell Signaling | 3743S |
| SEC61 | 1-1,000 | Cell Signaling | 14868 |
| cGAS | 1:300 | Cell Signaling | 15102 |

| | | | |
|---|---|---|---|
| GAPDH | 1-1,000 | Cell Signaling | 2118 |
| 53BP1 | 1-1,000 | Santa Cruz | 22760 |
| p53 | 1-1,000 | Santa Cruz | 126 |
| γH2AX | 1-1,000 | Cell Signaling | 2577 |
| **Immunofluorescence** | | | |
| STING | 1:500 | Cell Signaling | 13647 |
| S366 p-STING | 1:1200 | Cell Signaling | 50907 |
| p65 | 1:1000 | Cell Signaling | 8242 |
| Ki67 | 1:700 | Cell Signaling | 9129S |

**Supplemental Table 2.** Oligo sequences used in the study.

| **Human Gene** | **Sequence (5' → 3')** |
|---|---|
| *IFNB* | F - GTCAGAGTGGAAATCCTAAG |
| | R - ACAGCATCTGCTGGTTGAAG |
| *IL1B* | F - TGCACGCTCCGGGACTCACA |
| | R - CATGGAGAACACCACTTGTTGCTCC |
| *IL6* | F - CCTTCCAAAGATGGCTGAAA |
| | R - TTTCACCAGGCAAGTCTCCT |
| *IL8* | F - ACATGACTTCCAAGCTGGCC |
| | R - CAGAAATCAGGAAGGCTGCC |
| *STAT1* | F - ATCAGGCTCAGTCGGGGAATA |
| | R - TGGTCTCGTGTTCTCTGTTCT |
| *ISG15* | F - CGCAGATCACCCAGAAGATCG |
| | R - TTCGTCGCATTTGTCCACCA |
| *MX1* | F - GTTTCCGAAGTGGACATCGCA |

| | |
|---|---|
| | R - CTGCACAGGTTGTTCTCAGC |
| *IFIT3* | F - GAACATGCTGACCAAGCAGA |
| | R - CAGTTGTGTCCACCCTTCCT |
| *TMEM173* | F - ACTACTCCCTCCCAAATGCG |
| | R - GGCCACGTTGAAATTCCCTT |
| *CDKN2A* | F - ATCATCAGTCACCGAAGGTC |
| | R - CTCAAGAGAAGCCAGTAACC |
| *ACTB* | F - TGTACGCCAACACAGTGCTG |
| | R - GCTGGAAGGTGGACAGCGA |
| *GAPDH* | F - GCATGGCCTTCGGTGTCC |
| | R - AATGCCAGCCCCAGCGTCAAA |
| *ISG15* | F - CGCAGATCACCCAGAAGATCG |
| | R - TTCGTCGCATTTGTCCACCA |
| *cGAS* | F - ATGCAAAGGAAGGAAATGGT |
| | R - TTTAAACAATCTTTCCTGCAACA |
| *TMEM173* | sgRNA - TGAGTCACCTGGAGTGGATG |

**Supplemental Table 3.** Reagent and resources in this study.

| **Reagent** | **Company** | **Catalog number** |
|---|---|---|
| ADP | Milipore | 117105 |
| Baricitinib (JAK-STAT inhibitor) | TargetMol | T2485 |
| Blebbistatin | Cayman Chemical | 13013 |
| BSA | Sigma-Aldrich | A7906 |
| Chow diet | LabDiet | 5L0B |
| Creatine | Sigma-Aldrich | C0780 |
| DAPI | Vectashield | H-1800 |
| DMSO | Sigma-Aldrich | D2650 |
| Doxycycline | Sigma-Aldrich | D9891 |
| Dulbecco's modified Eagle's medium (DMEM) | Sigma-Aldrich | D5796 |
| FCCP | Cayman Chemical | 15218 |
| Fetal Bovine Serum (FBS) | Atlas Biologicals | EF-0500-A |
| Formaldehyde solution | Sigma-Aldrich | F8775 |
| Gel cream hair remover | Veet | 8329150 |
| Glucose | Agilent | 103577-100 |
| Glutamate | Sigma-Aldrich | G1251 |
| Glutamine | Agilent | 103579-100 |
| Octanoyl Carnitine | TOCRIS | 605 |
| Polyethylene glycol, PEG 300, | Sigma-Aldrich | 90878 |
| Power SYBR Green PCR Master Mix | Applied Biosystems | 4367659 |
| Pyruvate | Agilent | 103578-100 |
| Saponin | Sigma-Aldrich | 57900 |

| | | |
|---|---|---|
| Sodium pyruvate | Sigma-Aldrich | P2256 |
| Succinate | Sigma-Aldrich | S3674 |
| Triton X-100 | Fisher | BP151-500 |
| Tween 20 | Fisher | BP337500 |
| Tween 80 | Sigma-Aldrich | P1754 |
| UREA buffer | Sigma-Aldrich | 51457 |
